# Positive, negative and engineered durotaxis

**DOI:** 10.1101/2022.09.26.508682

**Authors:** P. Sáez, C. Venturini

**Affiliations:** Laboratori de Càlcul Numèric (LaCaN), Universitat Politècnica de Catalunya, Barcelona, Spain; E.T.S. de Ingeniería de Caminos, Universitat Politècnica de Catalunya, Barcelona, Spain; Institut de Matemàtiques de la UPC-BarcelonaTech (IMTech), Universitat Politècnica de Catalunya, Barcelona, Spain

**Keywords:** Durotaxis, Cell adhesion, Active Gels, Clutch Models, Tissue engineering

## Abstract

Cell migration is a physical process central to life. Among others, it regulates embryogenesis, tissue regeneration and tumor growth. Therefore, understanding and controlling cell migration represent fundamental challenges in science. Specifically, the ability of cells to follow stiffness gradients, known as durotaxis, is ubiquitous across most cell types. Even so, certain cells follow positive stiffness gradients while others move along negative gradients. How the physical mechanisms involved in cell migration works to enable a wide range of durotactic responses is still poorly understood. Here, we provide a mechanistic rationale of durotaxis by integrating stochastic clutch models for cell adhesion with an active gel theory of cell migration. We show that positive and negative durotaxis found across cell types are explained by asymmetries in the cell adhesion dynamics. We rationalize durotaxis by an asymmetric mechanotransduction in the cell adhesion behavior that further polarizes the intracellular retrograde flow and the protruding velocity at the cell membrane. Our theoretical framework confirms previous experimental observations and explains positive and negative durotaxis. Moreover, we show how durotaxis can be engineer to manipulate cell migration, which has important implications in biology, medicine and bioengineering.

## 1 Introduction

Cell migration is central to life [13, 29, 51]. It determines fundamental biological processes such as embryonic development, tissue regeneration, wound healing or tumor invasion. Cells move guided by exogenous chemical [50, 21], electrical [30, 7] and mechanical signals in-vivo and in-vitro [45, 49, 6, 51], known as chemotaxis, electrotaxis and durotaxis. During decades, there have been tremendous efforts to understand how cells organize themselves to follow these stimuli. There is also an increasing interest in controlling cell migration through external cues because it may allows us to propose strategies to arrest tumor progression [14, 57, 4], boost tissues regeneration [13, 42] and to design biomimetic materials [27, 53].

Durotaxis [49, 11, 45] represents a universal mode of directed cell migration across cell types. Among others, fibroblasts [28] and smooth muscle cells [54] migrate toward positive gradients of the the extracellular matrix (ECM), while others, e.g. neurons [25] or cancer cells [47], migrate toward negative gradients. These are referred to as positive and negative durotaxis, respectively. A stiffness gradient in the ECM exposes cells to a differential rigidity that they sense, transduce and integrate into intracellular responses. Experimental evidences showed dependences on the stiffness gradient [18, 25, 48] and on the absolute stiffness value in which cells migrate [8, 48]. However, there is a lack of physical understanding of durotaxis across cell scales that could help us to rationalize durotaxis. Among other insights, this could allow us to understand why some cells express positive durotaxis while others migrate down the stiffness gradient. Recent experimental work, in combination with clutch models, have shown the first evidences of how cell adhesion is implicated in these opposed durotactic responses [19].

As for any cell migration mode [56, 3], durotaxis should be driven by a competition of a continuous polymerization of actin filaments that protrudes the cell membrane forward [12, 44, 55] and an inward retrograde flow generated by the contractile forces that myosin motors exert on the F-actin network [35, 37] (Fig. 1). Balancing these two forces, adhesion complexes (ACs) establish cell attachements to their surroundings through a large number of cell adhesion molecules (CAMs) [36, 52]. At some point during the motility process, the formation of a cell front orients the cell toward positive or negative stiffness gradients.

**Figure 1:**
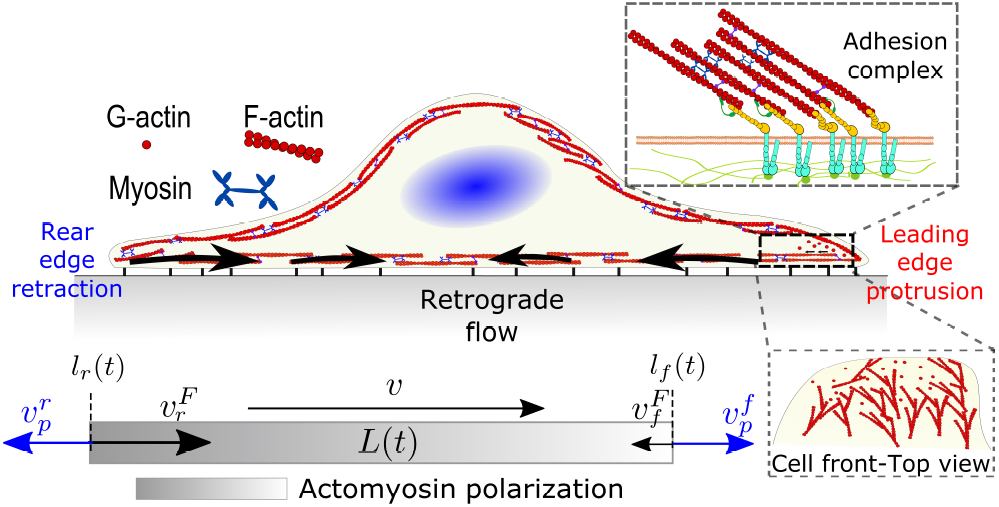
Sketch of the main forces acting in cell migration. A contractile acto-myosin flow, (black), competes with protrusive forces at the leading (red) and rear (blue) edges of the cell. At the cell-ECM interface, a friction-like force appears as a result of adhesion mechanisms.

Here, we integrate active gel models with stochastic clutch models for cell adhesion to reveal how durotaxis works. Cells read differences in the ECM stiffness through CAMs and regulate the ACs dynamics accordingly [38, 22, 52, 36]. Therefore, durotaxis must be first activated by the mechanotransduction of the substrate stiffness. Then, the striking durotactic modes found across cell types must be due to a differential expression of adhesion dynamics. Under this rationale, we explore how the cell adhesion can be manipulated to arrest or enhance durotaxis, and to switch from positive to negative durotaxis, and vice-versa.

## 2 Model

### 2.1 Clutch model of cell adhesion

To model the adhesion dynamics, we adopt previous clutch models (see [5, 15, 10] for details). The clutch hypothesis considers a contractile acto-myosin network that pulls on the CAMs bound to the ECM. Different cell types respond to the intracellular pulling forces in different ways because cells express specific types of CAMs. Some cells, e.g. neurons [5], express slip bonds: the life-time of the bond decreases exponentially as the force increases. Other cells, such as fibroblasts, express catch bonds, that is the lifetime first increases and then decreases exponentially with force. To consider these differences and the specific adhesion dynamics, the clutch model considers that a number of molecular clutches binds to the ECM with a constant rate *k*_*on*_, and unbinds with a dissociation rate 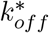, that depends on force as

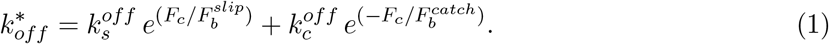

The first term in Eq. 1 alone, represents a slip bond. 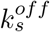 and 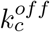 and 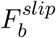 and 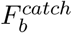 are the unloaded dissociation rates and the characteristic bond rupture forces in the slip and catch pathways respectively. *F*_*c*_ is force per molecular clucth. Some cells also show a reinforcement mechanism when the talin rod unfolds and vinculin binds to the unfolded domains, which promotes the increase of integrin density. Consequently, the traction forces that the cell exerts on the ECM rises [10].

The clutch model predicts the actin velocity, 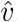, and tractions, *P*, as a function of the ECM stiffness (Fig. 2). We focus here on cells expressing slip bonds, as a case of non-reinforced bonds, and cells expressing reinforced catch bonds (all model parameters are in Table SI).

**Figure 2:**
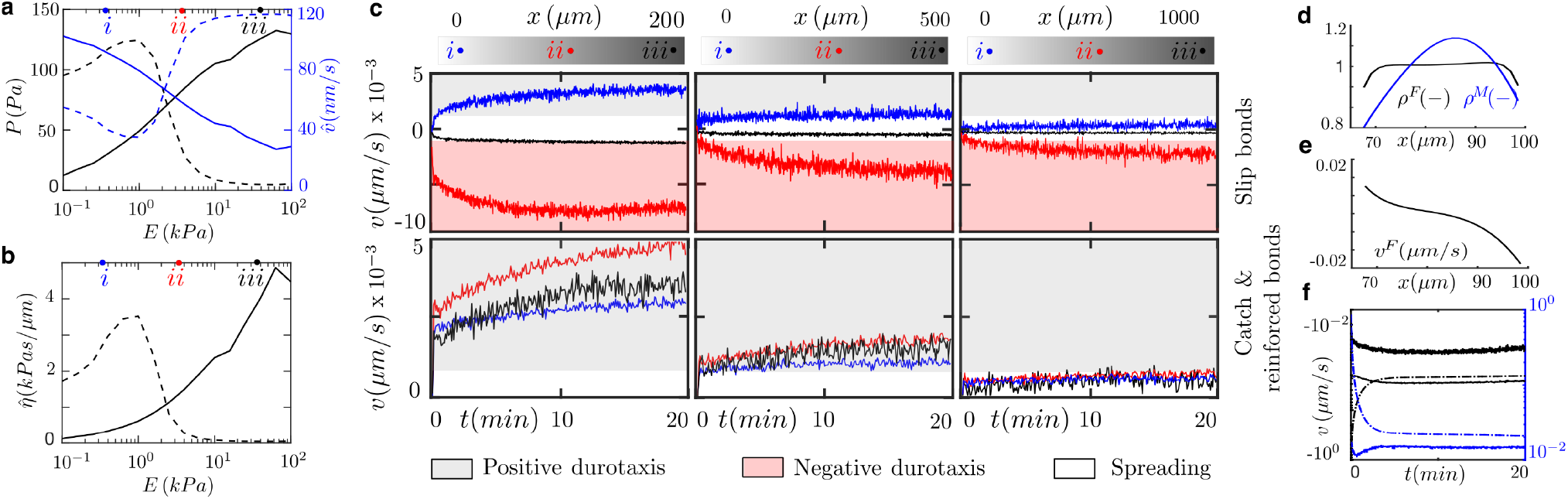
Results of computational clutch models for cells expressing slip (slip case) and reinforced catch bonds (reinforced case) for values of the substrate rigidity in 0.1-100 kPa. (a) Traction (black) and velocity (blue) for the slip case (dash) and reinforced case (solid). (b) Effective friction for the slip case (dash) and reinforced case (solid). Similarly to the tractions, friction increases monotonically for increasing values of the substrate rigidity in the reinforced case, while it presents a hill shape with a maximum at ≈ 1 kPa in the slip case, where the optimal rigidity localizes. (c) Migration velocities for cells expressing slip bonds (top) and catch bonds with talin reinforcement (bottom). Cells are followed for 20 min. The substrate stiffness goes from 0.1 kPa, at the left of the sample, to 100 kPa, at the right. Points i (blue), ii (red) and iii (black) represent the locations where the cells are initially seeded. In the sample of 200 *µ*m in length, i=33 *µ*m, ii=100 *µ*m, iii=166 *µ*m. In the sample of 500 *µ*m in length, i=83 *µ*m, ii=250 *µ*m, iii=416 *µ*m. In the sample of 1000 *µ*m in length, i=150 *µ*m, ii=500 *µ*m, iii=850 *µ*m. Grey shadow shows regions of positive durotaxis, red shadow represents negative durotaxis and white represents regions of symmetric spreading. (d-f) Singles cell durotaxis in a sample of 200 *µ*m in length for cells expressing slip bonds and seeded at E ≈ 3 kPa. At steady state, actin (black) and myosin (blue) densities (d) and retrograde flow in the lab (black) and cell (blue) frame (e). Time evolution of polymerization velocity (dot-dash), retrograde velocity at the cell membrane (dash) and total velocity of protrusion (solid) (f).

### 2.2 Minimal active gel model for cell migration

To explore the role of the adhesion mechanics in cell migration, we build a minimal active gel model for cell migration [41]. We consider a 1D domain, Ω, with moving coordinates *x*(*t*) ∈ [*l*_*r*_(*t*), *l*_*f*_ (*t*)]. *l*_*r*_(*t*) and *l*_*f*_ (*t*) represents the rear and front boundaries of the cell and, therefore, the cell length is determined as *L*(*t*) = *l*_*f*_ (*t*) − *l*_*r*_(*t*) (Fig. 1), where *f* and *r* indicate the front and rear of the cell.

We model the mechanics of the acto-myosin network as a contractile viscous gel, that represents the retrograde flow, in contact with the ECM [40, 43, 41, 31, 1]. We assume that viscous forces dominate the elastic forces and that inertial forces are negligible. Therefore, the balance of linear momentum for the active gel is

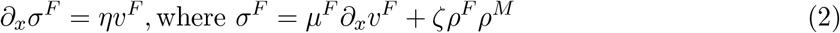

is the constitutive relation of the network stress, which accounts for the viscosity of the actin network and the myosin contractility, respectively. The right hand side of the balance of linear momentum represents the friction between the network flow and the ECM, with friction parameter *η. v*^*F*^ is the velocity of the retrograde flow, *µ*^*F*^ is the shear viscosity, *ζ* the active contraction exerted by the myosin motors and *ρ*^*M*^ and *ρ*^*F*^ are the densities of myosin motors and F-actin, respectively (see details below). We impose zero stresses on the cell boundaries so that there are not pushing or pulling forces on the network, which allows us to compute the resulting retrograde flow at the cell boundaries.

A second differentiated actin network polymerizes against the membrane [39, 37], which increases the membrane tension and, in turn, reduces the polymerization velocity [12, 44]. Filaments grow freely with velocity 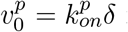 if there is no opposing force to it [32]. 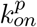 is the rate of actin assembly and *δ* is the size of one single monomer at the tip of the filament. When the membrane tension increases, the actin polymerization decreases as 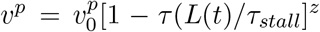 [32, 26, 23], where *τ*_*stall*_ is a tension required to stall the polymerization of the actin network and *z* is a model parameter that controls the velocity decay. We assume that actin polymerization is symmetric and *v*^*p*^ is equal at both sides of the cell. The membrane tension, *τ*, follows a simple Hookean law such that *τ* (*L*(*t*)) = *k*(*L*(*t*) − *L*_*b*_), where *k* is the spring constant and *L*_*b*_ = *L*_0_ + *L*_*r*_ accounts for the resting length, *L*_0_, and the buffer membrane length *L*_*r*_ in reservoirs and foldings of the cell membrane. Therefore, membrane tension starts to increase once all reservoirs and membrane folds have flatten, i.e *L*(*t*) *> L*_*b*_. We assume zero compressive stresses when *L*(*t*) *< L*_*b*_.

The outward polymerization velocities, 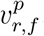, compete against the cell membrane with the inward retrograde flow velocity 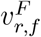 to expand or retract the cell rear and front of the cell with velocity 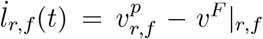, respectively (Fig. 1). Then, we compute the cell migration velocity as 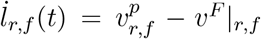.

The distribution of the F-actin density, *ρ*^*F*^ (*x, t*), is modeled as

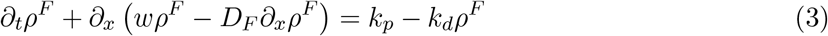

where the right hand side includes the polymerization and depolymerization terms with rates *k*_*p*_ and *k*_*d*_, respectively. *D*_*F*_ is the diffusive parameter of the F-actin. We write the transport equations in the cell frame and, accordingly, we define the velocity of the F-actin network as *w* = *v*^*F*^ − *v*. We impose zero fluxes on the cell boundaries to reflect that no F-actin can enter or leave the cell domain. Similarly, we model the myosin motors bound to the F-actin network [43, 2, 41], *ρ*_*M*_, as

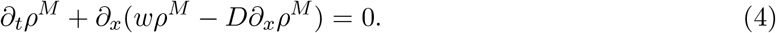

*D* is the effective diffusion parameter [2, 41]. The bound myosin also obeys zero flux boundary conditions to describe that no bound myosin motors can enter or leave the cell. All model parameters are summarized in Table SII.

### 2.3 Coupling the clutch and the active gel model

The friction between the actin network flow has two components, 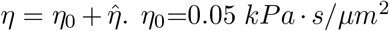 is a baseline friction of the retrograde flow with the surrounding cytoskeletal structures. 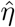 is an effective friction parameter 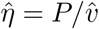 that arises from the friction created by the adhesion complexes. For cells expressing pure slip and catch bonds, there is a maximum traction force at a stiffness value, E≈ 1kPa, called optimum stiffness (Fig. 2a). Below and above the optimum stiffness, the traction forces increase and decrease with an increasing value of the substrate rigidity, respectively. However, reinforced bonds show that the traction forces increases as the stiffness of the ECM increases. For the slip case, the effective friction has also a maximum of ≈ 4 *kPa* · *s/µm*^2^ at the optimum rigidity (Fig. 2b). The friction is ≈ 2 *kPa* at 0.1 kPa and vanishes for substrate rigidities larger than 10 kPa. Friction increases monotically from zero to ≈ 4.5 *kPa* · *s/µm*^2^ in the reinforced case. Because ACs only exert tractions where they form, we weight these friction values by an averaged ACs density of 0.1, meaning that ACs occupies the 10% of the contact area. These differences in the cell adhesion behavior could explain the opposite durotactic modes found across cell types.

### 2.4 Numerical solution of the system of equations and model parameters

We solve computationally the system above (Material and Methods 2.2) in a staggered approach. We use the finite element method to discretize the system in space, and an implicit second-order Crank-Nicholson method to discretize the parabolic equations in time [60, **?**]. The numerical solution of the parabolic equations can present undesired oscillations if the problem becomes convective dominant, i.e. *Pe >* 1. The Peclet numbers for the actin and myosin transport problems are *Pe*_*F*_ = *hw/*2*D*_*F*_ and *Pe*_*M*_ = *hw/*2*D* respectively, where *h* is the size of the finite elements. As the convective velocity is the solution of the problem, we cannot guarantee a priori that the problem will remain in the limit case of *Pe <* 1. We use finite elements of constant size *h* = *l*_*f*_ (*t*)*/N*, where N is the number of elements. Because we want to keep the number of elements of our domain constant, and avoid remeshing strategies in the case that the element size needs to be decreased to keep *Pe <* 1, we include the Stream-Upwind Pretrov Galerkin (SUPG) stabilization term to overcome possible numerical oscillations in our solution [**?**]. The complete finite element and time discretization procedure is presented in Supplementary Material. The model parameters of the clutch and active gel model are summarized in Tables S1-S3.

## 3 Results

### 3.1 Cell adhesion explains positive and negative durotaxis

To analyze the exposure of cells to exogenous stimuli in the form of a shallow or a highly localized stiffness gradients, we use samples of varying length (200, 500 and 1000 *µ*m) with stiffnesses between 0.1 and 100kPa [10] (Fig. 2a). The clutch model predicts traction stresses and retrograde flow as a function of the ECM rigidity and bonds behaviour (see Fig. 2a and Materials and Methods). Then, an effective friction 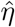 can be obtained to inform an active gel model for cell migration (see Materials and Methods) about the asymmetry that the stiffness gradient induce in the cell-ECM friction.

We initially seed cells at three different locations of the sample to study how the absolute value of the substrate stiffness, and not only its gradient, may regulate the durotactic response. We consider that cells undergo durotaxis if the migration velocity is larger than v=1 nm/s, otherwise we assume that they spread and remain stationary. We analyze the process during the first 20 minutes, when the durotactic response is already apparent.

Cells seeded in the sample of 1000 *µ*m in length do not express durotaxis. However, there is a strong durotactic response in cells seeded on the 200 *µ*m-length sample. Cells seeded on the 500 *µ*m-length sample also sustain durotaxis but at a lower degree. In the three samples (200, 500 and 1000 *µ*m), the strongest durotactic response is obtained for cells located at the center of the sample (E ≈ 1-10 kPa). This is due to a large friction gradient (Fig. 2b) and an absolute friction that enable the intracellular polarization.

In terms of migration direction (Fig. 2c), all cells with reinforced adhesions moves toward the positive stiffness gradient of the ECM with similar migration velocities that peak for substrate rigidities between 0.1-1 kPa. However, without reinforcement, cells can migrate toward positive or negative stiffness gradients of the ECM. Cells located on the left of the optimal rigidity (E = 0.1-1 kPa) undergo positive durotaxis. Cells located at the stiffest region of the samples, above 10 kPa, and where no friction gradient exists, show a weak durotactic response and stay mostly stationary. However, cells located just on the right of the optimal rigidity, E ≈ 1-10 kPa, express negative durotaxis. This is because of a negative friction gradient. For the strongest durotactic expression, in the 200 *µ*m-length sample and for cells seeded on the right of the optimal rigidity, the maximum migration velocity is *v* ≈ 8 nm/s. Cells seeded on the left of the optimal rigidity migrate with *v* ≈ 4.5 nm/s. However, cells seeded on E ≈ 10-100 kPa show a weak durotactic response.

To understand the physical mechanisms behind durotaxis, we further analyze the model results for the strongest durotactic response (right of the optimal rigidity, 200 *µ*m sample) of cells expressing slip bonds (Fig. 2a,c). We analyze the model during 20 min, when the migration velocity is maximum. The actomyosin network polarizes (Fig. 2d) because of an asymmetric retrograde flow (Fig. 2e) that results from asymmetric adhesion forces. The asymmetric retrograde flow and the constant polymerization velocity at both cell edges makes one side of the cell to protrude faster than the other, which establishes the cell front (Fig. 2f). Cells elongates 4-fold from the initial length and the cell membrane reaches a tension of ≈ 60 *pN/nm* (Fig. S1), similar to previous experimental data [46]. The stress of the actomyosin network is mostly symmetric with a maximum at the cell center of 50 *Pa* (Fig. S1). This result indicates that the actomyosin network imposes a traction on the nuclear region of the cell, which is also in agreement with previous results on the mechanosensitivity of the cell nucleus [9]. All other cases of durotaxis shown in Fig. 2 exhibit similar results, with the model variables polarized towards the positive or negative direction, depending on the direction of the friction polarization, and a smaller or larger polarization depending on the strength of the durotaxis response (Fig. S2). All cases with spreading signatures (see Fig. 2) show an almost symmetric distribution of all models variables (Fig. S3).

At steady state (Fig. 3), the friction values are symmetric and large enough to significantly reduce the retrograde flow, which reduces the front-rear differences between the inward flow and the outward actin polymerization velocity and, consequently, cells get into an stationary spreading-like phase.

**Figure 3:**
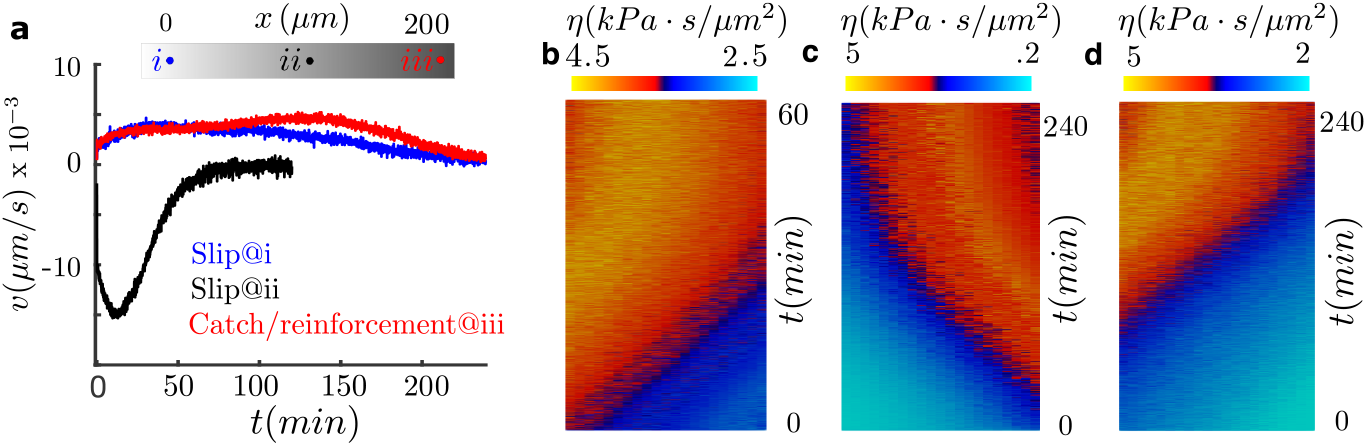
Analysis of durotaxis until steady state. All cells are in a sample of 200 *µ*m in length. For the slip cases (b-c), cells are located at soft (a, point i) and intermediate stiffness (a, point ii) locations. For the reinforced case, cells are seeded at stiff locations (a, point iii). (a) Migration velocity of the cell. (b-d) Kymographs of the cell friction for the three durotactic cases analyzed.

To further analyse the durotaxis dynamics as cells travel along the sample until they reach a steady state (reached when cells stop and no longer migrate), we focus on the three cases (Fig. S3): the two slip cases of cells initially seeded on the left and right of the optimum rigidity in the 200 *µ*m-length sample (strong positive and negative durotaxis expression, respectively, Fig. 2c) and the reinforced case with cells seeded on the stiff side, E ≈ 30 kPa, of the 200 *µ*m-length sample (positive durotaxis expression, Fig. 2c). As cells migrate, they travel through regions of different stiffnesses (Fig. S3b-d) and, therefore, the friction computed through the clutch model also changes in space and time (Fig. 3b-d). In the slip cases, cells migrate toward the optimal rigidity location and stop when they reach it (Fig. 3b and Fig. 3a). Cells stall at the optimal rigidity because they reach a symmetric distribution of the adhesion forces, which reverses the durotactic mode into the symmetric spreading phase (Fig. 3a-c). The non-motile steady state is reached at ≈ 1 and 4 hrs for cells expressing negative and positive durotaxis, respectively (Fig. 3). For the reinforced case, where no optimal rigidity exists, cells stall when they reach a friction of ≈ 4.5 *kPa* · *s/µm*^2^ at a stiffness of ≈ 60 kPa.

### 3.2 Durotaxis response to talin plasmids: switching to negative durotaxis

Then, we look into the effect of talin knockdown, because we know that it changes the cell adhesion behavior remarkably [10] and, ask ourselves if it could reverse positive durotaxis. By talin depletion, talin reinforcement is cancelled and, therefore, the adhesion behaves as a pure catch bond. This inhibition of the adhesion reinforcement induces a drastic reduction in cell traction and, consequently, in cell friction above a new optimal rigidity that forms at E≈7 kPa (Fig. 4a).

**Figure 4:**
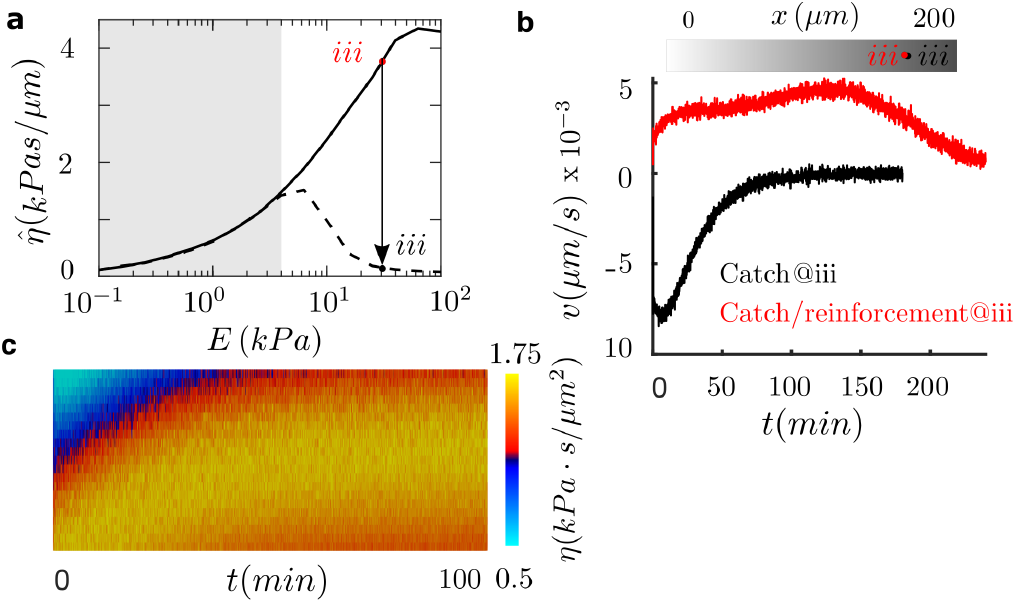
(a) Results of computational clutch models for cells expressing catch (catch case) and reinforced catch bonds (reinforced case) for values of the substrate rigidity between 0.1-100 kPa. Effective friction for the catch case (dash) and reinforced case (solid). Similarly to the traction forces, friction increases monotonically for increasing values of the substrate rigidity in the reinforced case, while in the catch case it presents a hill shape with a maximum at ≈ 1*kPa*. (b-c) Durotaxis for catch and reinforced cases until steady-state. All cells are in a sample of 200 *µ*m in length. Cells are located at the stiff side of the sample (a, point iii), where their behaviors differ. (b) Migration velocities along time and (c) kymographs of the cell friction for the reinforced case.

To analyze this idea, we use again a sample of 200 *µ*m in length and we seed cells at E ≈ 30 kPa. We choose this location to analyze substrate regions where the catch and the talin reinforced cases differ (white region, Fig. 4a). Our results show that we are able to shift the direction of durotaxis by inhibiting adhesion reinforcement (Fig. 4b). Furthermore, the motility of the cell is enhanced, and the maximum migration velocity doubles in the knockdown case to ≈ 8*nm/s*. These cells reach the target optimal rigidity in ≈ 90 min (Fig. 4b). At the optimal rigidity, the cell reaches a state of symmetric friction, which makes it to depolarize and stall. Therefore, the talin knockdown allows us to switch the directed durotaxis. Moreover, we could tune the location where the motility of the cell stalls and, therefore, we could design matrices to precisely control the target locations in cell migration.

### 3.3 Engineering durotaxis

Finally, we further explore possibilities for engineering durotaxis other than the matrix features where cells are seeded. Specifically, we are interested in physical quantities that may enhance, arrest or shift durotaxis. Our results above indicate that cell adhesion, in particular the asymmetric distribution of the adhesion strength, controls durotaxis. Therefore, we should arrest durotaxis when the traction force vanishes, are too high or there is no friction gradients, enhance it when the friction gradient increases, in combination with placing cells in specific stiffnesses, and shift the durotactic direction if the friction gradient changes sign. We focus on adhesion parameters that can clearly induce these changes in the adhesion behaviour and that are prone to be manipulated in-vitro and in-vivo. To do so, we performed a parametric analysis of different adhesion models parameters (the binding rate *k*_*ont*_, the slip part of the off-rate in all the unbinding models *k*_*off,slip*_, the catch part of the unbinding rate, *k*_*off,catch*_, the number of integrins added to the system *int*_*add*_ and the rate of vinculin attachment *k*_*onv*_). We show in Fig. 5 the specific values of the model parameters and the resulting traction.

**Figure 5:**
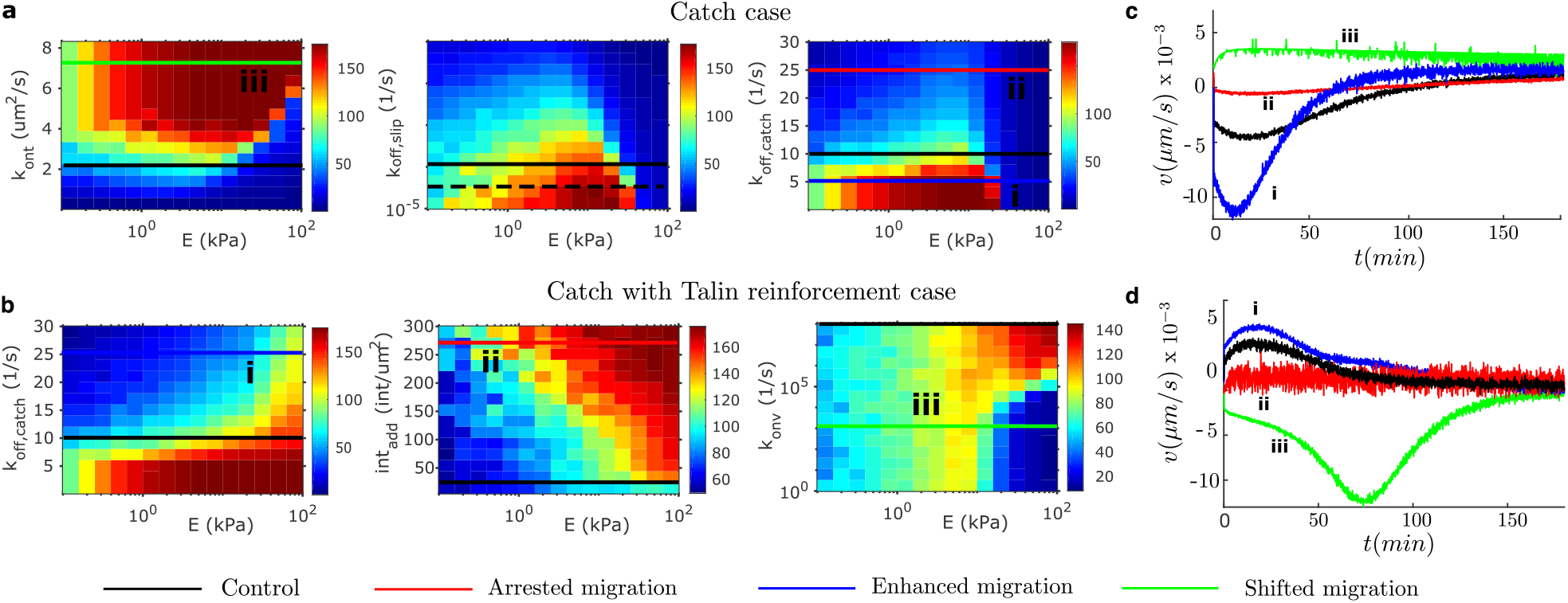
Results of computational clutch models for the traction force of cells expressing (a) pure catch bonds (catch case) and (b) reinforced catch bonds (reinforced case) for values of the substrate rigidity in 0.1-100 kPa. All cells are in a sample of 200 *µ*m in length. The control cases are plotted in black. We show cases of arrested (red, point ii) and enhanced (blue, point i) migration, and shifted migration direction (green, point iii). The the binding rate *k*_*ont*_ and the slip part of the off-rate in all the unbinding models *k*_*off,slip*_, the catch part of the unbinding rate *k*_*off,catch*_, the number of integrins added to the system *int*_*add*_, and the rate of vinculin attachment *k*_*onv*_ are analyzed. Migration velocity during durotaxis simulated until steady-state for catch (c) and reinforced cases of cell adhesion (d).

Among the different possibilities prone to change durotaxis, we present one case of enhancement, arrest and direction shifting of durotaxis for each adhesion case. We show that in the pure catch case, where the maximum migration velocity in the control case is ≈ 5 nm/s, we can enhance durotaxis by decreasing the off-rate of the catch bond (Fig. 5a,c). We show that the migration velocity increases to 12 nm/s (Fig. 5c). We also demonstrate a shift in the migration direction when the off-rate the integrin increases. These two modification are enable by increasing and decreasing the rigidity gradient, respectively. To shift the direction of migration, i.e. to shift the gradient sign, we increase the on binding rate of the integrin. Similarly, for the reinforced case (Fig. 5b), we can foster durotaxis by increasing the off-rate of the catch part of the bond. In this case, the migration velocity increases from ≈ 5 nm/s in the control case to ≈ 7 nm/s (Fig. 5d). We can arrest durotaxis by increasing the recruitment of integrins to the AC, which creates friction large enough to significantly reduce the retrograde flow. Finally, we can also shift the durotaxis direction, with a remarkable enhancement in the migration velocity of ≈ 12 nm/s (Fig. 5d), by reducing the on rate of vinculin to talin. This inhibition of vinculin binding cancels the adhesion reinforcement and, at the same time, increase the friction gradient (see also Fig. 4).

## 4 Discussion

Durotaxis represents a universal mode of directed cell migration. Although it manifests across most cell types, fibroblasts [28] and smooth muscle cells [54] express positive durotaxis, while others, e.g. neurons [25] and human fibrosarcoma cells [47], present negative durotaxis. How the physical mechanisms that enable cell motility cooperate to activate these remarkable and reversed modes of durotaxis were hindered behind complex in-vivo and in-vitro tests.

Different theoretical and computational models have explained positive and negative durotaxis individually (see, e.g, [33, 24, 34]). Recently, a clutch model in cooperation with experimental data showed the role of adhesion mechanics in dictating the durotactic direction. Here, we couple clutch models with classical active gel models for cell migration, from where we can recover actin flows and densities at a time and spatial scale. Our theory also explains positive and negative durotaxis by the mechanics of cell adhesion. Our results suggest that cells that express bonds with talin reinforcement follow positive durotaxis while those that express negative durotaxis are crowded by either pure slip or catch bonds. This is the case, e.g., of fibroblasts [10, 28] and neurons [5], respectively. In other words, reinforcement seems to control the durotactic response.

Our results also support and validate previous experimental data. We show that cells need a strong stiffness gradient in order to express durotaxis and that they prefer intermediate stiffnesses rather than soft or stiff substrates [18, 25, 48, 20, 16, 18]. Indeed, there is a sweet spot at 1-5 kPa that favors durotaxis. Conversely, lower and higher stiffnesses reduce cell motility, which are aligned with low and high friction forces, respectively, which can even arrest durotaxis. These results may also explain differences between weakly and strongly adherent cancer cells [58]. Interestingly, our results also show that cells migrate through the extracellular space guided by the stiffness gradient up to regions of large and symmetric adhesion forces, where the asymmetry of the motile forces stall and cells stop. These results indicate that cells migrate towards the optimal rigidity location, when it exists, or up to a location of a friction large enough to stall the polarization of the cell.

We also show specific aspects of the adhesion complex that can be pinpointed to control the strength and the direction of durotaxis. Future experimental work should confirm these predictions and further interact with computational models to exploit the control of cell migration. Because mechanical cues usually coexist with other exogenous signals, our theory can be extended to incorporate, e.g., chemotaxis and investigate the competition between these two prevalent migrating cues. Our theory explains durotaxis based on the polarized mechanical feedback from the ACs to the retrograde flow. However, a differential membrane tension could also induce indirect downstream signals to enhance actin and myosin activity and aid in the formation of a clear cell front and rear.

A deep understanding and precise control of the mechanisms in cell motility may allows us to not only understand how tumor cells invade healthy tissues and metastasize [14, 57] or how cells orchestrate the tissues regeneration [13, 42] but also provide tools to arrest tumor progression [4] and to engineer cell migration for better biomimetic tissue designs [17, 59].

## 5 Author Contributions

P.S designed the work, performed full simulations. C.V. performed the clutch model simulation.

P.S. wrote the paper. All authors analyzed the data and critically review the paper.

## 6 Acknowledgments

P.S acknowledges support from the the Spanish Ministry of Economy and Competitiveness (Grant No: PID2019-110949GB-I00) and the Generalitat de Catalunya (Grant No: 2017-SGR-1278). C.V. was supported by the Generalitat de Catalunya (FI grant). P.S and C.V. acknowledges support from the European Commission (Grant No. H2020-FETPROACT-01-2016-731957).

## References

[1] Erin Barnhart, Kun-Chun Chun Lee, Greg M. Allen, Julie A. Theriot, and Alex Mogilner. Balance between cell-substrate adhesion and myosin contraction determines the frequency of motility initiation in fish keratocytes. Proc. Natl. Acad. Sci. U. S. A., 112(16):5045–5050, 2015.

[2] Erin L. Barnhart, Kun Chun Lee, Kinneret Keren, Alex Mogilner, and Julie A. Theriot. An adhesion-dependent switch between mechanisms that determine motile cell shape. PLoS Biol., 9(5), 2011.

[3] James E. Bear and Jason M. Haugh. Directed migration of mesenchymal cells: Where signaling and the cytoskeleton meet. Curr. Opin. Cell Biol., 30(1):74–82, 2014.

[4] Darci T Butcher, Tamara Alliston, and Valerie M Weaver. A tense situation: forcing tumour progression. Nat Rev Cancer, 9(2):108–122, feb 2009.

[5] C E Chan and D J Odde. Traction dynamics of filopodia on compliant substrates. Science (80-.)., 322(5908):1687–1691, 2008.

[6] Guillaume Charras and Erik Sahai. Physical influences of the extracellular environment on cell migration. Nat. Rev. Mol. Cell Biol., 15(12):813–824, 2014.

[7] Barbara Cortese, Ilaria Elena Palamà, Stefania D’Amone, and Giuseppe Gigli. Influence of electrotaxis on cell behaviour. Integr. Biol. (United Kingdom), 6(9):817–830, 2014.

[8] Brian J. DuChez, Andrew D. Doyle, Emilios K. Dimitriadis, and Kenneth M. Yamada. Durotaxis by Human Cancer Cells. Biophys. J., 116(4):670–683, 2019.

[9] Alberto Elosegui-Artola, Ion Andreu, Amy E.M. Beedle, Ainhoa Lezamiz, Marina Uroz, Anita J. Kosmalska, Roger Oria, Jenny Z. Kechagia, Palma Rico-Lastres, Anabel Lise Le Roux, Catherine M. Shanahan, Xavier Trepat, Daniel Navajas, Sergi Garcia-Manyes, and Pere Roca-Cusachs. Force Triggers YAP Nuclear Entry by Regulating Transport across Nuclear Pores. Cell, 171(6):1397–1410.e14, 2017.

[10] Alberto Elosegui-Artola, Roger Oria, Yunfeng Chen, Anita Kosmalska, Carlos Pérez-gonzález, Natalia Castro, Cheng Zhu, Xavier Trepat, and Pere Roca-Cusachs. Mechanical regulation of a molecular clutch defines force transmission and transduction in response to matrix rigidity. Nat. Cell Biol., 18(October 2015):540, apr 2016.

[11] Jaime A. Espina, Cristian L. Marchant, and Elias H. Barriga. Durotaxis: the mechanical control of directed cell migration. FEBS J., 2021.

[12] Matthew J. Footer, Jacob W.J. Kerssemakers, Julie A. Theriot, and Marileen Dogterom. Direct measurement of force generation by actin filament polymerization using an optical trap. Proc. Natl. Acad. Sci. U. S. A., 104(7):2181–2186, 2007.

[13] Peter Friedl and Darren Gilmour. Collective cell migration in morphogenesis, regeneration and cancer. Nat. Rev. Mol. Cell Biol., 10(7):445–457, 2009.

[14] Peter Friedl and Katarina Wolf. Tumour-cell invasion and migration: diversity and escape mechanisms. Nat. Rev. Cancer, 3(5):362–374, 2003.

[15] Grégory Giannone, René Marc Mège, and Olivier Thoumine. Multi-level molecular clutches in motile cell processes. Trends Cell Biol., 19(9):475–486, 2009.

[16] William J. Hadden, Jennifer L. Young, Andrew W. Holle, Meg L. McFetridge, Du Yong Kim, Philip Wijesinghe, Hermes Taylor-Weiner, Jessica H. Wen, Andrew R. Lee, Karen Bieback, Ba Ngu Vo, David D. Sampson, Brendan F. Kennedy, Joachim P. Spatz, Adam J. Engler, and Yu Suk Cho. Stem cell migration and mechanotransduction on linear stiffness gradient hydrogels. Proc. Natl. Acad. Sci. U. S. A., 114(22):5647–5652, 2017.

[17] Donald E. Ingber, Van C. Mow, David Butler, Laura Niklason, Johnny Huard, Jeremy Mao, Ioannis Yannas, David Kaplan, and Gordana Vunjak-Novakovic. Tissue Engineering and Developmental Biology: Going Biomimetic. Tissue Eng., 12(12):3265–3283, 2006.

[18] Brett C. Isenberg, Paul A. DiMilla, Matthew Walker, Sooyoung Kim, and Joyce Y. Wong. Vascular smooth muscle cell durotaxis depends on substrate stiffness gradient strength. Biophys. J., 97(5):1313–1322, 2009.

[19] Aleksi Isomursu, Keun-Young Park, Jay Hou, Cheng Bo, Ghaidan Shamsan, Benjamin Fuller, Jesse Kasim, M Mohsen Mahmoodi, Tian Jian Lu, Guy M Genin, and Others. Negative durotaxis: cell movement toward softer environments. BioRxiv, 2020.

[20] Danielle Joaquin, Michael Grigola, Gubeum Kwon, Christopher Blasius, Yutao Han, Daniel Perlitz, Jing Jiang, Yvonne Ziegler, Ann Nardulli, and K Jimmy Hsia. Cell migration and organization in three-dimensional in vitro culture driven by stiffness gradient. Biotechnol. Bioeng., 113(11):2496–2506, 2016.

[21] Robert R. Kay, Paul Langridge, David Traynor, and Oliver Hoeller. Changing directions in the study of chemotaxis. Nat Rev Mol Cell Biol, 9(6):455–463, jun 2008.

[22] Jenny Z. Kechagia, Johanna Ivaska, and Pere Roca-Cusachs. Integrins as biomechanical sensors of the microenvironment. Nat. Rev. Mol. Cell Biol., 20(8):457–473, 2019.

[23] Kinneret Keren, Zachary Pincus, Greg M. Allen, Erin L. Barnhart, Gerard Marriott, Alex Mogilner, and Julie A. Theriot. Mechanism of shape determination in motile cells. Nature, 453(7194):475–480, 2008.

[24] Youjoung Kim, Seth M. Meade, Keying Chen, He Feng, Jacob Rayyan, Allison Hess-Dunning, and Evon S. Ereifej. Nano-architectural approaches for improved intracortical interface technologies. Front. Neurosci., 12(JUL):1–20, 2018.

[25] David E Koser, Amelia J Thompson, Sarah K Foster, Asha Dwivedy, Eva K Pillai, Graham K Sheridan, Hanno Svoboda, Matheus Viana, Luciano da F Costa, Jochen Guck, Christine E Holt, Kristian Franze, Luciano da F Costa, Jochen Guck, and Others. Mechanosensing is critical for axon growth in the developing brain. Nat. Neurosci., 19(12):1592–1598, 2016.

[26] Michael M. Kozlov and Alex Mogilner. Model of polarization and bistability of cell fragments. Biophys. J., 93(11):3811–3819, 2007.

[27] Robert Langer and David A. Tirrell. Designing materials for biology and medicine. Nature, 428(6982):487–492, 2004.

[28] C M Lo, H B Wang, M Dembo, and Y L Wang. Cell movement is guided by the rigidity of the substrate. Biophys. J., 79(1):144–152, 2000.

[29] Roberto Mayor and Sandrine Etienne-Manneville. The front and rear of collective cell migration. Nat. Publ. Gr., 17(2):97–109, 2016.

[30] Colin D McCaig, Ann M Rajnicek, Bing Song, and Min Zhao. Controlling cell behavior electrically: current views and future potential. Physiol. Rev., 85(3):943–978, jul 2005.

[31] Alex Mogilner and Angelika Manhart. Intracellular Fluid Mechanics: Coupling Cytoplasmic Flow with Active Cytoskeletal Gel. Annu. Rev. Fluid Mech., 50(1):347–370, 2018.

[32] Alex Mogilner and George Oster. Force generation by actin polymerization II: The elastic ratchet and tethered filaments. Biophys. J., 84(3):1591–1605, 2003.

[33] Elizaveta A. Novikova, Matthew Raab, Dennis E. Discher, and Cornelis Storm. Persistence-Driven Durotaxis: Generic, Directed Motility in Rigidity Gradients. Physical Review Letters, 118(7):1–5, 2017.

[34] Hadrien Oliveri, Kristian Franze, and Alain Goriely. Theory for Durotactic Axon Guidance. Phys. Rev. Lett., 126(11):118101, 2021.

[35] D Pantaloni, C Le Clainche, and M F Carlier. Mechanism of actin-based motility. Science (80-.)., 292(May):1502–1506, 2001.

[36] J. Thomas Parsons, Alan Rick Horwitz, and Martin A. Schwartz. Cell adhesion: integrating cytoskeletal dynamics and cellular tension. Nat. Rev. Mol. Cell Biol., 11(9):633–43, sep 2010.

[37] Thomas D. Pollard and Gary G. Borisy. Cellular motility driven by assembly and disassembly of actin filaments, 2003.

[38] Bruno Pontes, Pascale Monzo, Laurent Gole, Anabel Lise Le Roux, Anita Joanna Kosmalska, Zhi Yang Tam, Weiwei Luo, Sophie Kan, Virgile Viasnoff, Pere Roca-Cusachs, Lisa Tucker-Kellogg, and Nils C. Gauthier. Membrane tension controls adhesion positioning at the leading edge of cells. J. Cell Biol., 216(9):2959–2977, 2017.

[39] A. Ponti, M Machacek, S L Gupton, C M Waterman-Storer, and G Danuser. Two Distinct Actin Networks Drive the Protrusion of Migrating Cells. Science (80-.)., 305(5691):1782–1786, 2004.

[40] J. Prost, F. Jülicher, and J. F. Joanny. Active gel physics. Nat. Phys., 11(2):111–117, 2015.

[41] T. Putelat, P. Recho, and L. Truskinovsky. Mechanical stress as a regulator of cell motility. Phys. Rev. E, 97(1):1–11, 2018.

[42] Feini Qu, Farshid Guilak, and Robert L Mauck. Cell migration: implications for repair and regeneration in joint disease. Nat. Rev. Rheumatol., 15(3):167–179, 2019.

[43] Boris Rubinstein, Maxime F. Fournier, Ken Jacobson, Alexander B. Verkhovsky, and Alex Mogilner. Actin-myosin viscoelastic flow in the keratocyte lamellipod. Biophys. J., 97(7):1853–1863, 2009.

[44] Christian H. Schreiber, Murray Stewart, and Thomas Duke. Simulation of cell motility that reproduces the force-velocity relationship. Proc. Natl. Acad. Sci. U. S. A., 107(20):9141–9146, 2010.

[45] Adam Shellard and Roberto Mayor. Durotaxis: The Hard Path from In Vitro to In Vivo. Dev. Cell, 56(2):227–239, 2021.

[46] Zheng Shi, Zachary T. Graber, Tobias Baumgart, Howard A. Stone, and Adam E. Cohen. Cell Membranes Resist Flow. Cell, 175(7):1769–1779.e13, 2018.

[47] Samir P. Singh, Michael P. Schwartz, Justin Y. Lee, Benjamin D. Fairbanks, and Kristi S. Anseth. A peptide functionalized poly(ethylene glycol) (PEG) hydrogel for investigating the influence of biochemical and biophysical matrix properties on tumor cell migration. Biomater. Sci., 2(7):1024–1034, 2014.

[48] Raimon Sunyer, Vito Conte, Jorge Escribano, Alberto Elosegui-Artola, Anna Labernadie, Léo Valon, Daniel Navajas, José Manuel García-Aznar, José J Muñoz, Pere Roca-Cusachs, and Xavier Trepat. Collective cell durotaxis emerges from long-range intercellular force transmission. Science (80-.)., 353(6304):1157–1161, 2016.

[49] Raimon Sunyer and Xavier Trepat. Durotaxis. Curr. Biol., 30(9):R383–R387, 2020.

[50] Peter J M Van Haastert and Peter N Devreotes. Chemotaxis: signalling the way forward. Nat. Rev. Mol. Cell Biol., 5(8):626–634, 2004.

[51] Sjoerd Van Helvert, Cornelis Storm, and Peter Friedl. Mechanoreciprocity in cell migration. Nature Cell Biology, 20(1):8–20, 2018.

[52] M Vicente-Manzanares, C K Choi, and A R Horwitz. Integrins in cell migration – the actin connection. J. Cell Sci., 1473:199–206, 2009.

[53] Ulrike G. K. Wegst, Hao Bai, Eduardo Saiz, Antoni P. Tomsia, and Robert O. Ritchie. Bioinspired structural materials. Nat. Mater., 14(1):23–36, 2014.

[54] Joyce Y Wong, Alan Velasco, Padmavathy Rajagopalan, and Quynh Pham. Directed Movement of Vascular Smooth Muscle Cells on Gradient-Compliant Hydrogels. Langmuir, 19(5):1908–1913, mar 2003.

[55] Stephanie Wong, Wei Hui Guo, and Yu Li Wang. Fibroblasts probe substrate rigidity with filopodia extensions before occupying an area. Proc. Natl. Acad. Sci. U. S. A., 111(48):17176–17181, 2014.

[56] Kenneth M. Yamada and Michael Sixt. Mechanisms of 3D cell migration. Nat. Rev. Mol. Cell Biol., 20(12):738–752, 2019.

[57] Hideki Yamaguchi, Jeffrey Wyckoff, and John Condeelis. Cell migration in tumors. Curr. Opin. Cell Biol., 17(5 SPEC. ISS.):559–564, 2005.

[58] Benjamin Yeoman, Gabriel Shatkin, Pranjali Beri, Afsheen Banisadr, Parag Katira, and Adam J Engler. Adhesion strength and contractility enable metastatic cells to become adurotactic. Cell Reports, 34(10), mar 2021.

[59] Boyang Zhang, Anastasia Korolj, Benjamin Fook Lun Lai, and Milica Radisic. Advances in organ-on-a-chip engineering. Nat. Rev. Mater., 3(8):257–278, 2018.

[60] O C Zienkiewicz and R. L. Taylor. The Finite Element Method (Fluid Dynamics). Methods, 3:347, 2000.

