## Supplementary Information for "Positive, negative and engineered durotaxis"

### 1 Supplementary figures

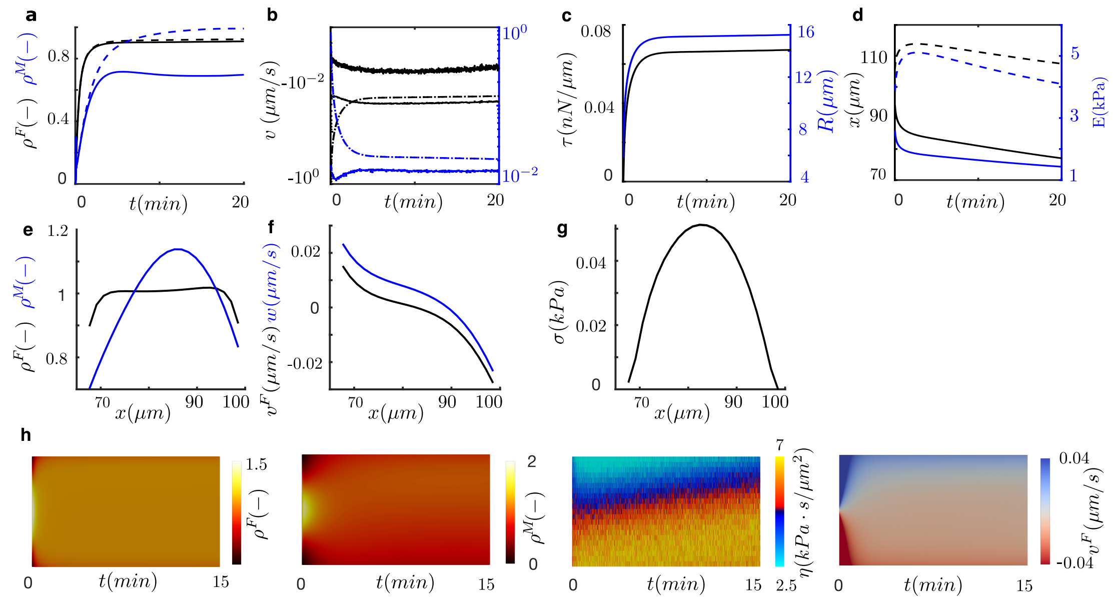

Figure 1: Negative durotaxis of single cells in a sample of 200 $\mu m$ in length for cell expressing slip bonds and seeded at E $\approx$ 3 kPa. (a) Actin (black) and myosin (blue) densities at the front (dash) and rear (solid) of the cell. (b) At the front and rear of the cell: polymerization velocity (dot-dash) and retrograde velocity at the cell membrane (dash). Total velocity of protrusion (solid). Negative velocities are in black, positive velocities are in blue. (c) Membrane tension (black) and cell radius (blue). (d) Position (black) of the cell rear (solid) and front (dash) and stiffness (blue) seen by the cell front and rear. At steady-state, (e) actin (black) and myosin (blue) densities, (f) retrograde flow in the lab (black) and cell (blue) frame and (g) tension of the actomyosin network. (h) Kymographs of the actin density, myosin density, adhesion friction and retrograde flow velocity.

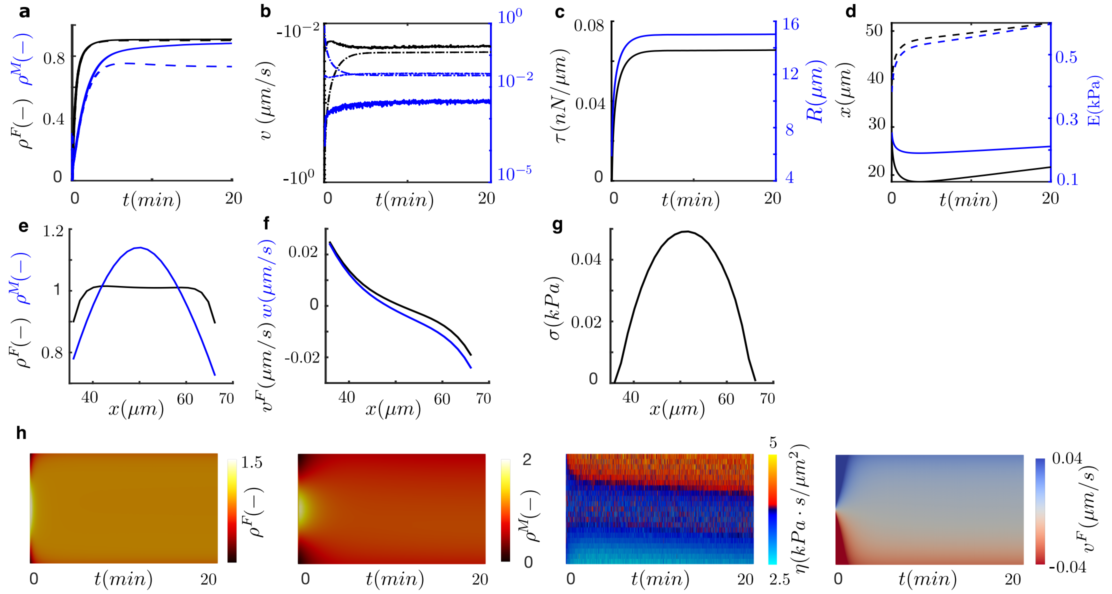


Figure 2: Positive durotaxis of single cells in a sample of 200 $\mu m$ in length for cell expressing slip bonds and seeded at E $\approx$ 0.3 kPa. (a) Actin (black) and myosin (blue) densities at the rear (dash) and front (solid) of the cell. (b) At the front and rear of the cell: polymerization velocity (dot-dash) and retrograde velocity at the cell membrane (dash). Total velocity of protrusion (solid). Negative velocities are in black, positive velocities are in blue. (c) Membrane tension (black) and cell radius (blue). (d) Position (black) of the cell front (solid) and rear (dash) and stiffness (blue) seen by the cell front and rear. At steady-state, (e) actin (black) and myosin (blue) densities, (f) retrograde flow in the lab (black) and cell (blue) frame and (g) tension of the actomyosin network. (h) Kymographs of the actin density, myosin density, adhesion friction and retrograde flow velocity.

#### 2 Clutch model

The "motor-clutch" model [9] uses a stochastic approach to analyze the effect of substrate stiffness in the tractions exerted by the cell. This conceptual framework includes some of the most important mechanisms in cell adhesion, pulling forces of myosin motors inside the actin network, which induce an F-actin retrograde flow, and the binding and unbinding dynamics of adhesion clutches with the ECM.

Actin filaments flow from the cell leading edge where adhesions form towards the cell center. Inside an AC, the molecular clutches link the actin filaments with the ECM, and are allowed to bind and unbind. The molecular clutches are made of several adaptor proteins, among which the most important are vinculin, talin, integrin and fibronectin. Vinculin binds to talin. Talin links the actin filaments at the top, to the integrin at the bottom. Finally, the transmembrane protein integrin, connects to a ligand in the ECM, like fibronectin. Usually, only one type of bond

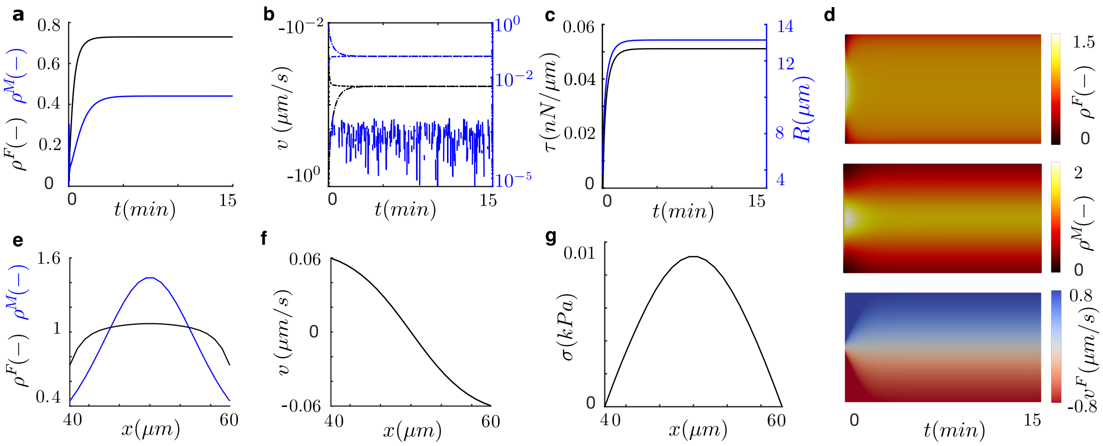


Figure 3: Time and space evolution of cell spreading. (a) Actin (black) and myosin (blue) densities at the left (dash) and right (solid) fronts of the cell. Left and right variables are superimposed as a result of their symmetric distributions. (b) At the front and rear of the cell: polymerization velocity (dot-dash) and retrograde velocity at the cell membrane (dash). Total velocity of protrusion (solid). Negative velocities are in black, positive velocities are in blue. (c) Membrane tension (black) and cell radius (blue). (d) Kymographs of the actin density (top), myosin density (center) and retrograde flow velocity (bottom). At steady-state, along the cell length, (e) actin (black) and myosin (blue) densities, (f) retrograde flow at the cell (blue) and lab frame (black) and (g) tension of the actin network.

| **Parameters** | | |
|---|---|---|
| $\mu^F$ [kPa·s] | 10 | [1, 2] |
| $\eta_0$ [kPa·s/$\mu$m] | 0.05 | [1, 3, 2] |
| $\zeta$ [kPa] | 0.05 | [1, 4] |
| $k$ | 0.1 | [2] |
| $D_F$ [$\mu$m/s] | 0.2 | [1, 5] |
| $D$ [$\mu$m/s] | 0.4 | [1, 5] |
| $k_d$ | 0.1 | [6, 7] |
| $k_p$ | 0.1 | [6, 7] |
| $\delta$ [nm] | 2.2 | [8] |
| $k_{on}$ [$s^{-1}$] | 250 | [8] |
| $L_0$ [$\mu$m] | 10 | * |
| $L_b$ [$\mu$m] | 0.3*$L_0$ | * |
| $\tau_{stall}$ [$nN/\mu$m] | 0.4 | * |

Table 1: Model parameters, values adopted and references to publications where they were obtained. * indicates values used in this work.

is considered, which is assumed to be the ”weakest link” in the adhesion chain. Bound binders deform and transmit force to the underlying substrate. Moreover, the stiffness of the substrate controls the force loading rate, i.e. the speed at which force is built in bound clutches, thus

| **Parameters** | **Slip** | **Catch** |
|---|---|---|
| | [9] | [14] |
| $E$ (kPa) | 0.1 - 100 | 0.1 - 100 |
| $a$ (nm) | 447 | 1700 |
| $n_c$ | 75 | 1200 |
| $\kappa_c$ (pN / nm) | 5 | 1000 |
| $k_{ont}$ ($\mu$m$^2$ / s) | 1 | 2.11e-4 |
| $d_{int}$ (int /$\mu$m$^2$) | 1 | 300 |
| $k_{off,slip}$ (s$^{-1}$) | 0.1 | - |
| $F_{b,slip}$ (pN) | 2 | - |
| $F_m$ (pN) | 2 | 2 |
| $n_m$ | 75 | 800 |
| $v_u$ (nm / s) | 120 | 110 |
| $k_{onv}$ kPas$^{-1}$) | - | 1e8 |
| $int_{add}$ (int /$\mu$m$^2$) | - | 24 |
| $m_r$ (int /$\mu$m$^2$) | - | 15000 |

Table S 2: Parameters adopted for the clutch model of cell adhesion. For the slip case, the model parameters are obtained from [9] and for the catch bonds with talin reinforcement are obtained from [14]. The unbinding rate is consider to follow a cycle mechanical reinforcement, following $k^*_{off} = 0.00079938\ e^{(F_c/8.16)} + 10.14\ e^{(-F_c/6.24)} + 900\ e^{(-F_c/0.01)}$ [14].

| **Parameters** | | |
|---|---|---|
| $\mu^F$ [kPa·s] | 10 | [1, 2] |
| $\eta_0$ [kPa·s/$\mu$m] | 0.05 | [1, 3, 2] |
| $\zeta$ [kPa] | 0.05 | [1, 4] |
| $k$ | 0.1 | [2] |
| $D_F$ [$\mu$m/s] | 0.2 | [1, 5] |
| $D$ [$\mu$m/s] | 0.4 | [1, 5] |
| $k_d$ | 0.1 | [6, 7] |
| $k_p$ | 0.1 | [6, 7] |
| $\delta$ [nm] | 2.2 | [8] |
| $k_{on}$ [$s^{-1}$] | 250 | [8] |
| $L_0$ [$\mu$m] | 10 | * |
| $L_b$ [$\mu$m] | 0.3*$L_0$ | * |
| $\tau_{stall}$ [$nN/\mu$m] | 0.4 | * |

Table S 3: Parameters adopted for the active gel model of cell migration and references to publications where they were obtained. * indicates values adjusted in this work.

influencing the resulting cell traction and retrograde flow. The model is computed through Monte Carlo simulations.

In short, the model is built as follow: Inside each MC simulation, at each time step, the molecular clutches are first allowed to bind to the F-actin bundle with binding rate $k_{on}$:

$$k_{on} = k_{ont} d_{int}, \tag{1}$$

where $k_{ont}$ is the constant binding rate and $d_{int}$ is the density of integrins on the membrane. Then, the bound clutches can unbind following their dissociation rate $k^*_{off}$. Binders following a slip behaviour have an unbinding rate which increases exponentially with force according to Bell's Model:

$$k^*_{off} = k_{off,slip} e^{\left(\dfrac{F_c}{F_{b,slip}}\right)}, \tag{2}$$

where $k_{off,slip}$ is the dissociation rate in absence of force, $F_{b,slip}$ is the characteristic bond rupture force, and $F_c$ is the force inside one binder:

$$F_{c(i)} = \kappa_c (x_{c(i)} - x_{sub}). \tag{3}$$

$\kappa_c$ is the stiffness of the clutch, $x_{c(i)}$ is the displacement of the molecular clutch $i$ and $x_{sub}$ is the displacement of the substrate. In the catch bonds, the $k^*_{off}$ becomes:

$$k^*_{off} = k_{off,slip} e^{\left(\dfrac{F_c}{F_{b,slip}}\right)} + k_{off,catch} e^{\left(\dfrac{-F_c}{F_{b,catch}}\right)}, \tag{4}$$

where $k_{off,slip}$ and $k_{off,catch}$ are the dissociation rates in absence of force via the slip and catch pathway respectively. $F_{b,slip}$ and $F_{b,catch}$ are the characteristic bond rupture forces via the slip and catch pathway, respectively. We will refer to the model of a cell expressing slip or catch bonds without talin reinforcement as slip and catch cases, respectively. All binders behave independently of the others, and the time at which one event happens is computed, e.g. for the binding event, as:

$$\tau_i = \frac{-\ln \xi_i}{k_{on(i)}}, \tag{5}$$

where $\xi_i$, $i = 1 \ldots n_c$, are independent random numbers uniformly distributed over $[0, 1]$, and $n_c$

is the total number of binders. The time of all possible events is computed and only the events happening before a fixed time step $\Delta t$ are executed.

After all the binding and unbinding events have been updated, the bound probability $P_b$ is calculated as:

$$P_b = \frac{n_{eng}}{n_c}, \tag{6}$$

where $n_{eng}$ is the number of engaged binders. The clutches and the substrate are treated as simple Hookean springs, i.e. the applied force scales linearly with respect to the displacement. Thus the force on the substrate $F_{sub}$ is:

$$F_{sub} = \kappa_{sub} x_{sub}. \tag{7}$$

The actin velocity $v_f$ is inversely related to the substrate force $F_{sub}$, as a loaded substrate will decrease the movement of the actin filaments. The force-velocity relation is given by:

$$v_f = v_u \left( 1 - \frac{F_{sub}}{F_{stall}} \right), \tag{8}$$

where $v_u$ is the unloaded velocity of myosin and $F_{stall}$ is the total myosin motors stall force, defined as:

$$F_{stall} = n_m F_m. \tag{9}$$

$F_m$ is the stall force of one myosin motor and $n_m$ is the number of motors. Once the state of the cluster with free and bound clutches is found, the F-actin velocity is calculated using the value of $F_{sub}$, and bound clutches are displaced by $\Delta x = v_f \Delta t$. The new displacements of all clutches $x_{c(i)}$ are computed, and the new displacement of the substrate becomes:

$$x_{sub} = \frac{\kappa_c \sum_{i=1}^{n_{eng}} x_{c(i)}}{\kappa_{sub} + n_{eng} \kappa_c}. \tag{10}$$

From the displacement of the substrate, the force against which the myosin motors are working, $F_{sub}$, is determined and is used to get the actin velocity for the following time step. Also the force along each molecular clutch is obtained and is used to compute the rate $k^*_{off}$ for the next step. The Young's modulus $E$ of the substrate is related to the substrate stiffness $\kappa_{sub}$ through the relation [10]:

$$\kappa_{sub} = \frac{E 4 \pi a}{9}, \tag{11}$$

where $a$ is the radius of the AC. Moreover, the cell traction $P$ applied to the substrate is computed as:

$$P = \frac{F_{sub}}{\pi a^2}, \tag{12}$$

where $\pi a^2$ is the area occupied by the $n_c$ molecular clutches considered in the AC.

The clutch model was extended in order to consider the effect of talin unfolding and vinculin binding in the adhesion behaviour [11]. Here, the binding and unbinding rates exactly refer to the integrin-fibronectin bond, which is considered to behave as a catch bond. Talin unfolding is a mechanosensing event triggered by force: the actin-integrin adaptor protein talin unfolds under force and exposes binding sites to vinculin. Talin unfolding follows a slip bond behaviour, i.e. when a force is applied, the unfolding time decreases exponentially with force. For low forces, integrin unbinding is faster than talin unfolding (the binder breaks and the force goes to zero), whereas for high forces, talin unfolding is faster. Therefore, above a stiffness threshold, talin unfolds revealing cryptic vinculin binding sites, and vinculin binds leading to integrin recruitment and adhesion reinforcement. Below that stiffness threshold, integrins unbind before talin can unfold, so no more integrins are recruited.As talin is not modelled as a protein with its proper spatial position, though its unfolding mechanics is taken into account, we consider the binder chain spatially made of substrate and integrin, directly bound to actin, so we define the displacement of the integrin as $x_{int(i)} = x_{c(i)} - x_{sub}$.

Now the binding rate $k_{on}$ in Eq. 1 evolves during time, as the density of integrins $d_{int}$ can change. At each time step, the unbinding rate $k^*_{off}(F)$ for the integrin-fibronectin catch bond and the unfolding rate $k_{unf}(F)$ for the talin slip bond, are computed for each bound clutch. Unbinding and unfolding times are determined stochastically according to $k^*_{off}$ and $k_{unf}$.If unfolding happens before unbinding, then either vinculin can bind to talin with a force-independent rate $k_{onv}$, or talin can refold with rate $k_{fold}(F)$.

If vinculin binding occurs, there is adhesion reinforcement and the integrin density, $d_{int}$, is increased by $int_{add}$ integrins/$\mu m^2$. If vinculin binding does not occur, then integrin density is decreased by $int_{add}$, reflecting that adhesions shrink if force application is decreased [12] [13]. However integrin density is never allowed to go below the initial value, nor above the maximum integrin density, $m_r$, as integrins cannot be closer than a minimum distance. We call $d_{int}^0$ the initial density of integrins. This model with talin reinforcement has been able to explain the experimental results in Mouse Embryonic Fibroblasts (MEFs) [14].

### References


[1] Larripa K, Mogilner A (2006) Transport of a 1D viscoelastic actin-myosin strip of gel as a model of a crawling cell. *Phys. A Stat. Mech. its Appl.* 372(1 SPEC. ISS.):113–123.

[2] Putelat T, Recho P, Truskinovsky L (2018) Mechanical stress as a regulator of cell motility. *Phys. Rev. E* 97(1):1–11.

[3] Bergert M, et al. (2015) Force transmission during adhesion-independent migration. *Nat. Cell Biol.* 17(4):524–9.

[4] Rubinstein B, Fournier MF, Jacobson K, Verkhovsky AB, Mogilner A (2009) Actin-myosin viscoelastic flow in the keratocyte lamellipod. *Biophys. J.* 97(7):1853–1863.

[5] Barnhart EL, Lee KC, Keren K, Mogilner A, Theriot JA (2011) An adhesion-dependent switch between mechanisms that determine motile cell shape. *PLoS Biol.* 9(5).

[6] Wilson CA, et al. (2010) Myosin II contributes to cell-scale actin network treadmilling through network disassembly. *Nature* 465(7296):373.

[7] Raz-Ben Aroush D, et al. (2017) Actin Turnover in Lamellipodial Fragments. *Curr. Biol.* 27(19):2963–2973.e14.

[8] Mogilner A, Oster G (2003) Force generation by actin polymerization II: The elastic ratchet and tethered filaments. *Biophys. J.* 84(3):1591–1605.

[9] Chan CE, Odde DJ (2008) Traction dynamics of filopodia on compliant substrates. *Science (80-. ).* 322(5908):1687–1691.

[10] Ghibaudo M, et al. (2008) Traction forces and rigidity sensing regulate cell functions. *Soft Matter* 4(9):1836–1843.

[11] Elosegui-Artola A, et al. (2014) Rigidity sensing and adaptation through regulation of integrin types. *Nat. Mater.* 13:631.

[12] Balaban NQ, et al. (2001) Force and focal adhesion assembly: a close relationship studied using elastic micropatterned substrates. *Nat. Cell Biol.* 3(5):466–472.

[13] Riveline D, et al. (2001) Focal Contacts as Mechanosensors: Externally Applied Local Mechanical Force Induces Growth of Focal Contacts by an Mdia1-Dependent and Rock-Independent Mechanism. *J. Cell Biol.* 153(6):1175–1186.

[14] Elosegui-Artola A, et al. (2016) Mechanical regulation of a molecular clutch defines force transmission and transduction in response to matrix rigidity. *Nat. Cell Biol.* 18(October 2015):540.
